# Complement facilitates developmental microglial pruning of astrocyte and vascular networks

**DOI:** 10.1101/2022.01.12.476001

**Authors:** Gopalan Gnanaguru, Steven J. Tabor, Gracia M. Bonilla, Ruslan Sadreyev, Kentaro Yuda, Jörg Köhl, Kip M. Connor

## Abstract

Microglia, the resident immune cell of the central nervous system (CNS), play a pivotal role in facilitating neurovascular development through mechanisms that are not fully understood. This current work resolves a previously unknown role for microglia in facilitating the developmental pruning of the astrocytic template resulting in a spatially organized retinal superficial vascular bed. Mechanistically, our study identified that local microglial expression of complement (C)3 and C3aR is necessary for the regulation of astrocyte patterning and vascular growth during retinal development. Ablation of retinal microglia and deficiency of C3 or C3aR reduced developmental pruning and clearance of astrocytic bodies leading to increased astrocyte density, altered vascular patterning, and elevated gene expression levels of extracellular matrix proteins involved in vascular growth during retinal development. These data demonstrate that C3/C3aR signaling is an important checkpoint required for the finetuning of vascular density during neuroretinal development.

## Introduction

High-energy demanding neural tissues, such as the retina, require the development of a complex vascular network for their growth, survival, and function. The process of vascular development in the neural retina is orchestrated through highly regulated multicellular crosstalk for the establishment of functional blood vessels [1–3]. In this process, the resident immune cell of the CNS, microglia, play a vital role in promoting vascular development [4, 5]. However; the mechanism(s) through which microglial cells dictate vascular growth in the retina is not fully resolved. Here, we explored the cellular and molecular contributions of microglia in the formation of intricate and spatially organized astrocyte and retinal vascular networks.

The development of the vascular system in the retina is mediated by astrocytes that enter the retina through the optic nerve head around birth in mice [6–8]. Astrocyte migration and template formation in the retina is an essential step for vascular development, and interestingly, species that lack astrocytes in the retina remain avascular [9,7,10]. In the vascularized retina, ganglion cell-derived Pdgfa acts as a chemoattractant that stimulates astrocyte entry into the retina at the time of birth in mice [6, 11]. As astrocytes proliferate and migrate, they form a spatially organized extracellular matrix enriched template that allows subsequent vascular growth to occur [12,6,13–15] . The leading endothelial tip cells follow the astrocyte laid template (beginning at P0) to form the primary superficial vascular plexus by P7 [12, 13], followed by the formation of an interconnected deep and intermediate vascular plexus that completes around P15 [2]. Disruption of astrocyte template formation perturbs the growth of subsequent primary and interconnected vascular plexus [16,14,17,15]. Although the importance of astrocyte migration into the retina for vascular growth is known, it remains unclear what signaling mechanism(s) modulate the spatial organization of astrocyte and vascular networks during retinal development. In cases of perturbed or abnormal vascular growth, permanent vision loss can occur [18–20], thus understanding these mechanism(s) of vascular development could lead to new strategies for treating retinal pathologies.

Microglial invasion into the neuroretina precedes astrocytes entry [21] and microglia are known to contribute to vascular growth by facilitating endothelial tip cell anastomoses [4, 5]. However, to date, it is less clear if microglia also interact with astrocytes to assist astrocytic template formation. During neural development, microglia play an indispensable role in the formation of a defined and functional neural circuitry mediated in part through the complement system [22–24]. The complement system is an integral component of the innate immune system, initially discovered as a component involved in opsonization and clearance of foreign pathogens [25]. Recent studies imply that in addition to its immune response function, the complement system regulates a wide-variety of biological functions such as synaptic pruning, regulation of glial-microglial crosstalk, clearance of abnormal neovascular growth, removal of stressed or dying cells, and modulation of cell metabolism [22,26–32,24,33,34]. Depending on the context/stimuli, complement activation is initiated through the classical, alternative, or lectin pathways [35], all of which lead to cleavage of C3 and C5 fragments, i.e. the large C3b and C5b and the small C3a and C5a fragments [25, 36]. Particularly, during neural development, the release of cleaved C3a binding and activation of its cognate C3aR regulates diverse biological functions such as the development of the cerebellum, modulation of embryonic neural progenitor proliferation, and chick eye morphogenesis [37,27,38]. While, C3b binding of CR3 (also known as integrin alpha M) facilitates microglial pruning of synapses [24]. This suggests a more ubiquitous role of C3 during retinal vascular development.

During superficial retinal vascularization, the astrocyte template undergoes rearrangement and a significant number of astrocytes are eliminated through an unknown mechanism [39, 40]. Here, we hypothesized and tested if C3 facilitates microglial pruning of the astrocytic template required for subsequent primary superficial vascular network formation. Our findings illustrate the importance of complement mediated pruning by microglia for the proper development of astrocytic and vascular networks.

## Results

### Microglia closely interact with astrocytes and endothelial cells during postnatal vascular development

Spatially organized honeycomb shaped matrices formed by astrocytes are critical for endothelial migration and formation of the primary superficial vasculature in the retina [12,6,14,15]. As microglia populate the retina at embryonic stages [21], we examined if microglia closely associate with astrocytes to facilitate template assembly for subsequent vascular growth. To distinguish microglia, we used the specific P2ry12 marker [41, 42]. P2ry12-labeled retinal microglia exhibited more prominent structures as compared to staining with Iba1 or isolectin B4 (**Fig. 1a, 1b and Supplementary Fig.1a**). Analysis of P0 retinal flatmounts revealed the close association of microglial ramified processes with the developing astrocytic template as well as the entering endothelial tip cells around the optic nerve head (ONH) region (**Fig. 1c**). In addition, at P0, we noticed the enveloping of microglial processes over astrocytes that are beginning to form a naïve template in the central and peripheral retinal regions required for subsequent vascular growth (**Fig. 1c**). Further developmental analysis at P5 illustrated the existence of intimate interactions between microglia, astrocytes, and endothelial tip cells in the vascularized regions and microglia-astrocyte interactions in the avascular regions (**Fig. 1d**).

**Figure 1.**
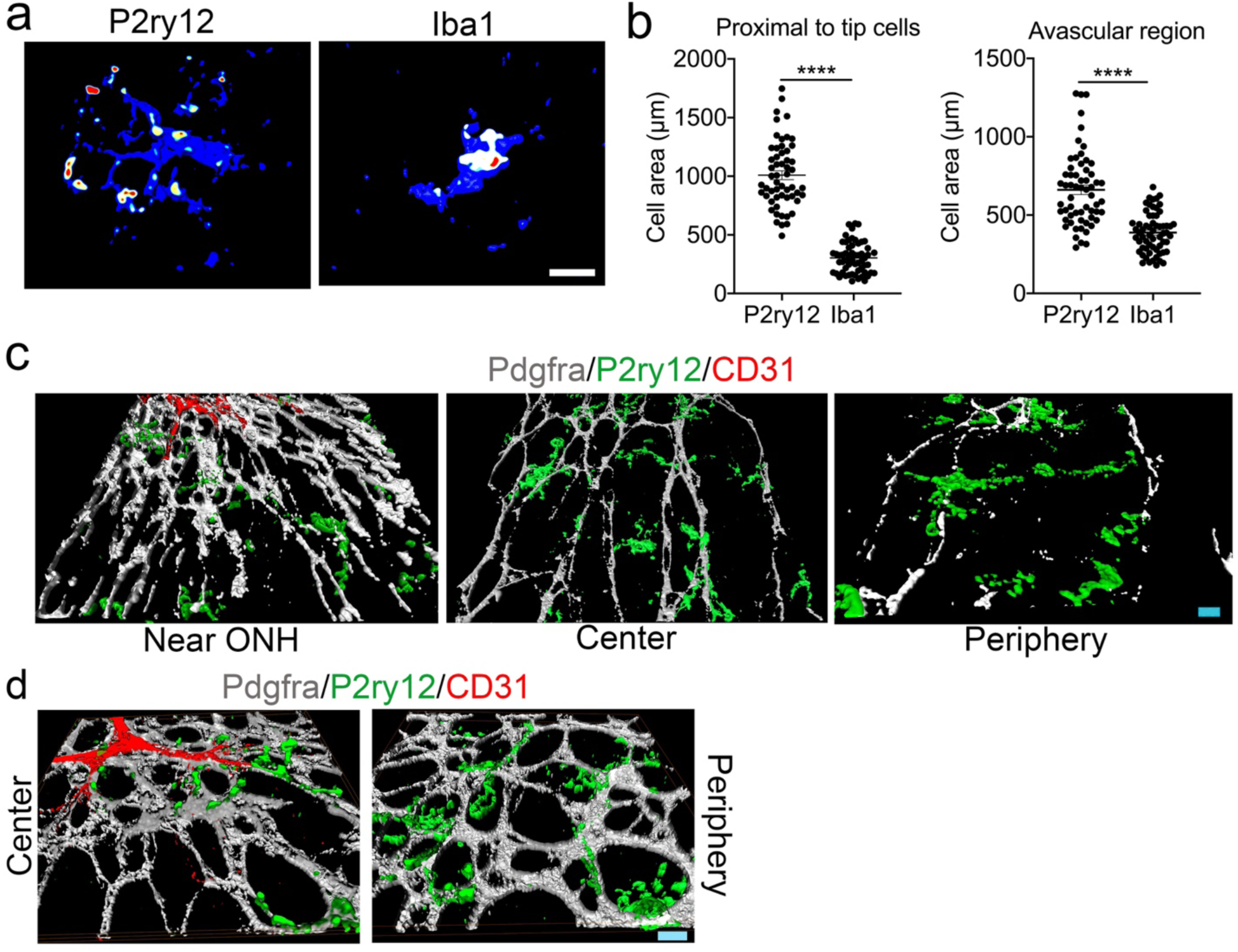
Microglia are recruited to sites of astrocyte template formation and vascularization. **a.** Representative 3D-reconstructed P5 retinal flatmounts immunostained for either P2ry12 or Iba1 revealing microglial morphology. Areas with increased intensity are indicated as hotspots (n=4). **b.** Quantification of P2ry12 and Iba1 localized expression around tip cells and avascular regions of the retina (n=4). (**c** and **d**) Retinal flatmounts prepared from P0 and P5 pups were immunostained for Pdgfra, P2ry12, and CD31. Z-stack images were then acquired and 3D-reconstructed to examine microglial interaction with astrocytes and endothelial tip cells early during vascular development. **c.** Representative 3D-reconstructed image of P0 C57BL/6J retinal flatmount near the optic nerve head (ONH), central retina, and peripheral retina revealing microglia (P2ry12-green) have a close association with astrocytes (Pdgfra-grey), and endothelial tip cells (CD31-red) (n=3). **d.** Representative 3D-reconstructed image of P5 C57BL/6J retinal flatmount near the tip cell area and avascular peripheral retina revealing microglial (P2ry12-green) have a close association with astrocytes (Pdgfra-grey), and endothelial tip cells (CD31-red) (n=5). Scale bars: **a**, **c**, and **d** are 10 µm. All error bars represent ± S.E.M. Statistical differences were calculated by unpaired *t*-test. **** P < 0.0001

### Depletion of microglia is associated with increased astrocytic density and spatial patterning

To further investigate the functional relevance of microglial interactions with astrocytes and endothelial cells during the critical steps of astrocytic template assembly and subsequent vascular development, we designed a strategy to deplete microglia from birth. Colony-stimulating factor 1 receptor (Csf1r) signaling is required for microglial survival [43,42,44], and prior work found that Csf1r deletion during early development is embryonic lethal [45], thus we utilized a pharmacologic approach whereby Csf1r is inhibited to deplete microglia [42, 44]. To overcome the technical challenges to deplete microglia from birth, timed-pregnant C57BL6J females were fed a diet incorporated with Csf1r specific antagonist (PLX5622) from E13.5/E14.5. We have recently reported that mice fed a diet containing Csf1r specific antagonist (PLX5622) effectively depletes microglia in the retina, similar to genetic approaches with minimal off-target effects [42, 44].

P2ry12 immunostaining of P1 and P5 retinal flatmounts of litters from control diet fed group or PLX5622 diet fed group showed that the PLX5622 antagonist successfully depleted microglia in comparison to controls (**Supplementary Fig. 1b and 1d)**. Analysis of RNA extracted from P1 and P5 retinas of PLX5622 group also showed significant decrease in Csf1r, Tmem119, and P2ry12 gene expression levels in comparison to the P1 control group (**Supplementary Fig. 1c and 1e**). Following microglial depletion during early retinal vascular development, we examined P1 (active astrocyte template assembly phase) and P5 (active astrocyte template rearrangement and vascular growth phase) retinas of litters from control mice and in mice lacking microglia.

Although microglial depletion did not prevent astrocyte migration into the retina at P1 (**data not shown**), examination of P5 retinal flatmounts from mice lacking microglia displayed abnormal clustering of astrocytes in the central vascular region and in the peripheral avascular region (**Fig. 2a and 2b**). While at P5, control retinal flatmounts displayed typical honeycomb patterning of astrocytes in the central vascular and peripheral avascular retina (**Fig. 2a**). Furthermore, quantitative analysis by flowcytometry showed that the percentage of Pdgfra+ (a marker for astrocytes) astrocyte population was significantly increased in pups lacking microglia in comparison to the control group (**Fig. 2d and 2e**). We also verified that PLX5622 treatment did not alter the expression level of retinal astrocytes growth factor *Pdgfa* [6, 11] (**Fig. 2c).** Intriguingly, the abnormal astrocytic density and clustering phenotype persisted even after the completion of vascular development at P15 (**Supplementary Fig. 3a-c**). Astrocyte distribution in P15 retinal flatmounts with and without microglia were assessed using the Gfap marker, as it is strongly expressed in mature astrocytes [39]. Gfap immunostained P15 retinas of pups lacking microglia showed a significant increase in astrocyte distribution along with increased astrocytic template branching in the central and peripheral retinas compared to controls (**Supplementary Fig. 3a and 3b**). P15 retinal flatmounts that lacked microglia also contained abnormal clustering of astrocytes compared to control retinas (**Supplementary Fig. 3c**). Taken together, these findings strongly implicate microglia in playing a significant role in supporting the spatial establishment of the astrocytic template required for retinal angiogenesis.

**Figure 2.**
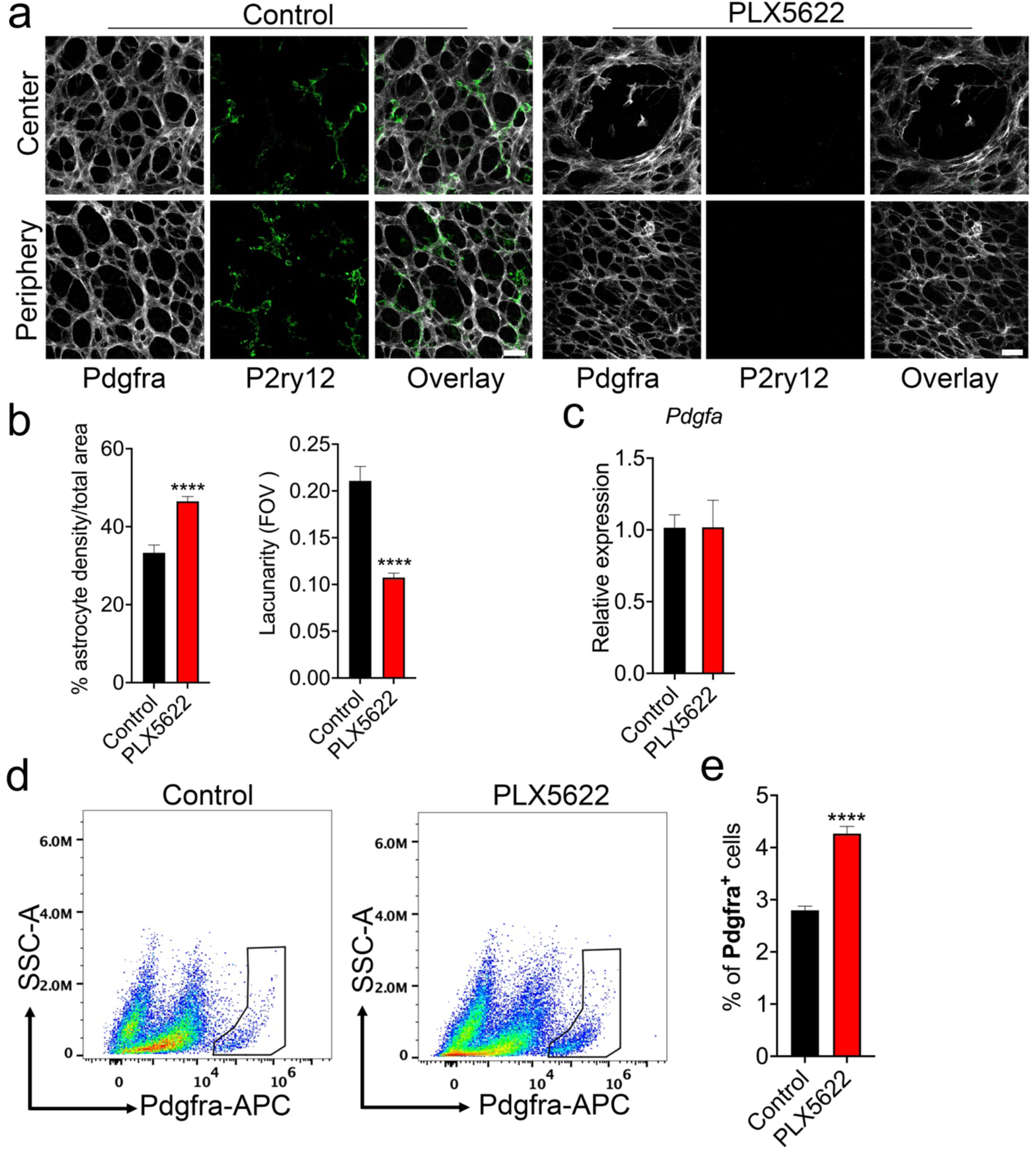
Microglial depletion in early vascular development increases astrocyte density and causes abnormal template formation. Time-pregnant C57BL/6J mice from gestational day E13.5/E14.5 were fed either a control or Csfr1 specific antagonist (PLX5622) diet and littermates were analyzed at P5. (a-e). **a.** Representative P5 retinal flatmounts from control or PLX5622 groups immunostained for Pdgfra (astrocytes) and P2ry12 (microglia) showing microglial distribution and patterning of astrocytes in the avascular mid/peripheral retina (n=4). Scale bar: 50 µm. **b.** Bar graph represents the ‘ImageJ AngioTool’ generated quantitative measurements of astrocyte density and lacunarity in the retinas of P5 control and PLX5622 groups immunostained for Pdgfra (n=5). **c.** Real-time PCR analysis showing *Pdgfa* gene expression levels at P5 in the retinas of control and PLX5622 groups (n=5)**. d.** Astrocyte (Pdgfra-APC+) live cell frequencies were assessed in P5 retinas of control and PLX5622 groups by flow cytometry (n=3). **e.** Quantification of flow cytometry data showing the percentage of Pdgfra-APC+ live cells in control and PLX5622 groups. All error bars represent ± S.E.M. Statistical differences between control and PLX5622 group were calculated by unpaired *t*-test **** P < 0.0001.

### Depletion of microglia reduced subsequent astrocyte dependent vascular growth and density

Proper formation of the astrocytic template and microglia-derived signaling are necessary for endothelial tip cell migration and subsequent formation of the superficial retinal vascular network [16,14,46,47,17,15]. As microglial depletion resulted in defective assembly of the astrocytic template, we next examined the effects on subsequent vascular growth. Analysis of retinal flatmounts in litters from mice lacking microglia showed that the vascularized area at P1 and P5 were significantly reduced in comparison to litters from the control group (**Supplementary Fig. 2a and 2b**). Depletion of microglia also significantly reduced vascular density and branching points, while dramatically increasing lacunarity in P5 litters of mice lacking microglia compared to litters from the control group (**Supplementary Fig. 2c and 2d**). We also verified that the PLX5622 diet did not have any off-target effects on suppressing Vegf isoforms expression levels (**Supplementary Fig. 1f**), which are required for retinal vascular growth [13,48,49]. There was an increase in the expression levels of the *Vegfa* isoform in microglia-depleted retinas compared to those of controls ((**Supplementary Fig. 1f**), and it could be due to increased astrocyte density [50]. Examination of P15 retinal flatmounts from mice lacking microglia revealed that the vascular phenotype observed (reduced density and increased lacunarity) persisted even after the completion of vascular development (**Supplementary Fig. 3d-f**). These results were consistent with the previous findings that microglia-derived proangiogenic factors are necessary for retinal vascular growth [4,46,5,47]. As microglial depletion altered spatial patterning of astrocytes (**Fig. 2**), we next focused on characterizing the role of microglia in regulating astrocyte patterning during superficial retinal vascular growth.

### Microglia utilize complement to facilitate spatial astrocytic template formation necessary for subsequent vascular development

During retinal vascular development, large numbers of astrocytes are eliminated through an uncharacterized mechanism likely facilitated by microglia peaking at P5 [39, 40], potentially to assist the establishment of the spatially organized astrocytic template. In agreement with previous findings [39, 40], analysis of P5 retinal flatmounts showed engulfed Gfap positive astrocytic cellular debris localized within the microglial endosomal/lysosomal membrane protein CD68 (**Supplementary Fig. 4a**).

C3 is well known to facilitate the refinement of neural architecture by pruning and eliminating dying neurons [22, 24]. Therefore, we investigated if C3 regulates patterning of astrocyte matrices through microglia for the establishment of a spatially organized vascular network. Assessment of P5 retinal flatmounts of C3 tdTomato reporter mice showed robust localization around the vascular growth front zone (**Fig. 3a**). High resolution images revealed C3 tdTomato expression in microglia that are closely associated with astrocytes in the vascular growth and adjacent avascular zones at the active astrocyte template rearrangement and subsequent vascular developmental phases (**Fig. 3b**).

**Figure 3.**
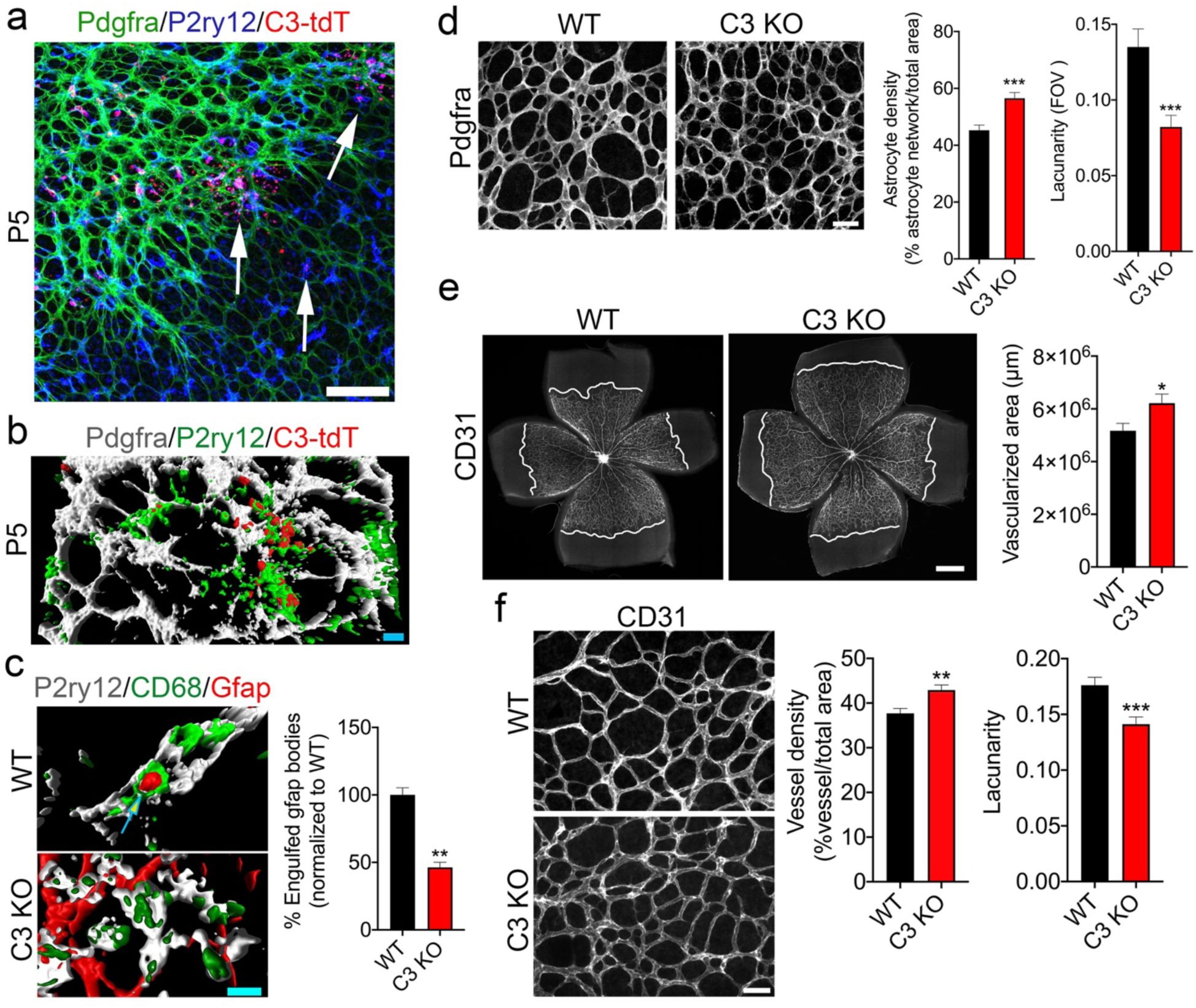
C3 is actively expressed at P5, and loss of C3 increases astrocyte and vascular density. **a.** Representative C3-tdTomato (C3-tdT) expressing P5 retinal flatmount immunostained for P2ry12 and Pdgfra to label microglia (blue) and astrocytes (green), respectively (n=5). **b**. Representative 3D-reconstructed image from P5 C3-tdT expressing retinal flatmount immunostained for P2ry12 and Pdgfra revealing C3 reporter expression within microglia (P2ry12, green) that are closely interacting with astrocytes (Pdgfra, grey) (n=5). **c**. Z-stack images were acquired from P5 WT and C3 KO retinal flatmounts immunostained for P2ry12, Gfap, and CD68 and then 3D-reconstructed to visualize microglia (P2ry12) and engulfed astrocyte (Gfap) bodies within microglial lysosomal/endosomal membrane protein CD68. Bar graph shows the percentage of Gfap debris localized within P2ry12, CD68 co-labelled microglia between WT and C3 KO retinas (n=3). **d.** Representative P5 retinal flatmounts from WT and C3 KO immunostained for Pdgfra showing astrocyte template patterning in the avascular region (n=5). Bar graphs represent the quantitative measurements of astrocyte density and lacunarity using ImageJ AngioTool (n=5). **e.** Representative P5 Retinal flatmounts of WT and C3 KO immunostained for CD31 showing vascular outgrowth and quantification of vascularized area in WT and C3 KO (n=5). **f.** Representative P5 retinal flatmounts of WT and C3 KO immunostained for CD31 displaying vascular density in the central retina. Differences in vessel density and spatial branching (lacunarity) were quantified in WT and C3 KO (n=5) using ‘ImageJ Angiotool’. All error bars represent ± S.E.M. Statistical differences between WT and C3 KO calculated by unpaired *t*-test * P < 0.05, ** P < 0.01, *** P < 0.001. Scale Bars: **a** is 75µm, **b** and **c** are 10 µm, **e** is 500µm, and **f** is 50µm.

We next assessed P5 retinal flatmounts of C3 deficient mice along with their WT littermate controls to define how the loss of C3 affected astrocytic template rearrangement. Analysis of C3 deficient P5 retinas showed a significant reduction in the localization of Gfap positive astrocytic cell debris within microglial CD68 compared to WT littermate control retinas (**Fig. 3c and Supplementary Fig. 4B**). Notably, in comparison to the WT P5 littermate retinal flatmounts, analysis of Pdgfra immunostained P5 C3 deficient retinal flatmounts displayed a significant increase in astrocyte template density and decreased lacunarity, indicative of reduced spatial gap in the astrocyte template (**Fig. 3d**). Intriguingly, analysis of P5 C1q KO retinal flatmounts showed no significant differences in astrocyte patterning or vascular growth (**Supplementary Fig. 5a and 5b**).

Given that the loss of C3 dramatically increased astrocyte density and spatial patterning (**Fig. 3d**), we next analyzed the effects of C3 loss on subsequent vascular growth at P5. In comparison to WT littermates, P5 C3 deficient retinal flatmounts showed a significant increase in the vascularized area (**Fig. 3e**). In addition to increased vascular growth, vessel density was significantly increased in the P5 retinal flatmounts of C3 deficient pups compared to P5 WT littermate retinal flatmounts (**Fig. 3f**), while the spatial gap was significantly reduced in P5 C3 KO compared to WT littermate controls (**Fig. 3f**). We then took a whole-genome RNA-seq approach to determine the molecular mechanism(s) for increased astrocyte and vascular density in C3 deficient retinas (**Fig 3**). Analysis of P5 C3 deficient retinas showed about 451 genes that were significantly differentially expressed in comparison to WT (**Fig. 4a and supplementary excel**). Consistent with findings thus far, gene expression profiling further confirmed that astrocyte specific markers, Gfap and Pax2, were significantly increased in C3 deficient P5 retinas compared to P5 WT retinas (**Fig. 4d**). Further pathway enrichment analysis of RNA-seq data using EnrichR [51] revealed an enrichment of genes linked to extracellular matrix (ECM) organization and biogenesis in C3 deficient P5 retinas (**Fig. 4b and 4c**). Notably, in P5 C3 deficient retinas, significantly elevated ECM genes included several genes involved in vascular development and integrity (**Fig 4d**) [52,14,53]. These results suggest that C3 plays an important regulatory role during the assembly of spatially organized astrocyte matrices and succeeding superficial retinal vascular growth.

**Figure 4.**
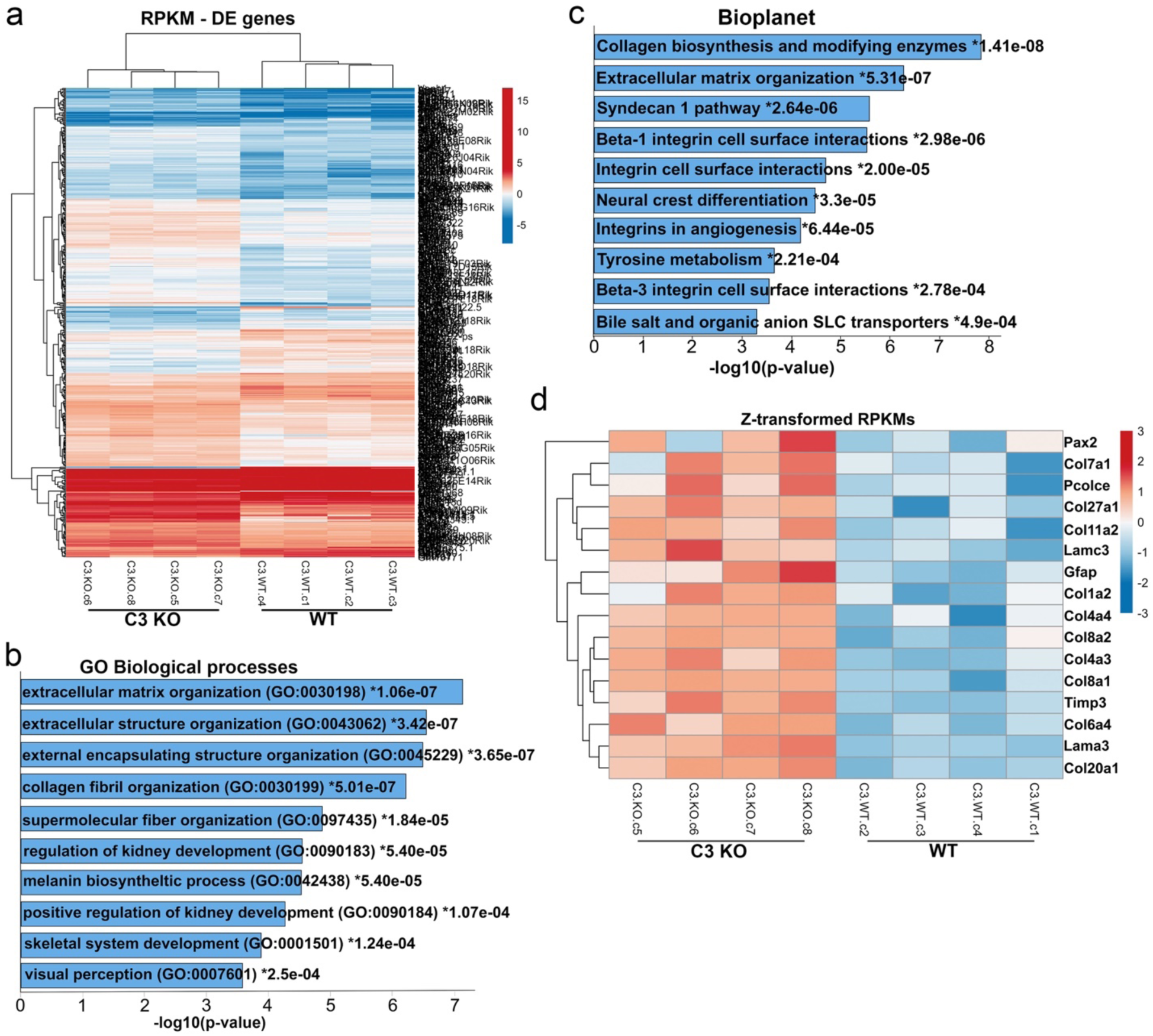
Loss of C3 increases transcripts levels of astrocyte markers and several extracellular matrix molecules involved in angiogenesis at P5. (**a-d**) Poly-A captured mRNA sequencing was performed on RNA extracted from P5 retinas of WT and C3 KO to determine differences in global gene expression patterns during the active superficial vascular development phase. **a.** Expression heatmap revealing global differential gene expression pattern in P5 WT and C3 KO retinas (n=4). **b.** Top ten Gene ontology (GO) biological processes that are upregulated in P5 retinas of C3 KO in comparison to WT according to EnrichR analysis. **c.** Top ten Bioplanet pathways that are upregulated in C3 KO retinas compared to WT retinas at P5 according to EnrichR analysis. **d.** Heatmap showing elevated gene expression levels (Z-score normalized RPKMs) of extracellular matrix proteins including astrocyte-specific Pax2 and Gfap transcripts in P5 C3 KO compared to WT retinal samples.

### C3aR controls microglia-mediated astrocytes template patterning

During retinal development, activation of CR3 by cleaved large C3b fragment is shown to facilitate microglial removal of ganglion cells [22]. Analysis of P5 CR3 KO retinal flatmounts did not show significant changes in astrocyte patterning or vascular growth (**Supplementary Fig. 5c and 5d**). As a recent study reported microglial phagocytosis of dying neurons through C3aR [54], we investigated if microglial C3aR-dependent mechanism is involved in the phagocytic elimination of astrocytes to create a spatially organized template for subsequent superficial vascular growth. Examination of P5 retinal flatmounts of C3aR-tdTomato reporter mice [55], showed strong expression of C3aR in microglia within vascular growth and avascular zones (**Fig. 5a and 5b**). We then tested if C3aR is involved in the phagocytic elimination of astrocyte bodies. We isolated and cultured retinal microglia and astrocytes from P5 pups. After labeling cultured astrocytes with DiI to track and identify this population, cell death was induced by staurosporine treatment. Retinal microglia were incubated with DiI-labeled dead astrocytic bodies in the presence or absence of C3aR neutralizing antibodies or control IgG. Blocking of C3aR significantly reduced microglial engulfment of DiI-labeled dead astrocytic bodies when compared to IgG control (**Fig. 5c**), revealing the phagocytic capability of microglial C3aR. Further examination of P5 C3aR KO retinal flatmounts showed a significant reduction in the localization of Gfap positive astrocytic cell debris within microglial CD68 compared to WT littermate control retinas (**Fig. 5d**). Analysis of Pdgfra+ astrocytic template formation at P5 also showed increased density and reduced spatial distribution in C3aR KO retinas compared to WT littermate controls (**Fig. 6b)**. We also evaluated the astrocyte specific *Pax2* gene expression and the expression of RNA-seq identified ECM genes involved in vascular development (Fig. 4d) in P5 C3aR KO and WT retinas. Data show that the expression levels of *Pax2*, *Col4a3*, and *Col8a1* were significantly increased in C3aR KO retinas similar to C3 deficient retinas (**Fig. 6a and 4d**), there was also an upward trend in *Lamc3, Col4a4*, and *Col8a2* genes expression levels compared to WT (**Fig. 6a**).

**Figure 5.**
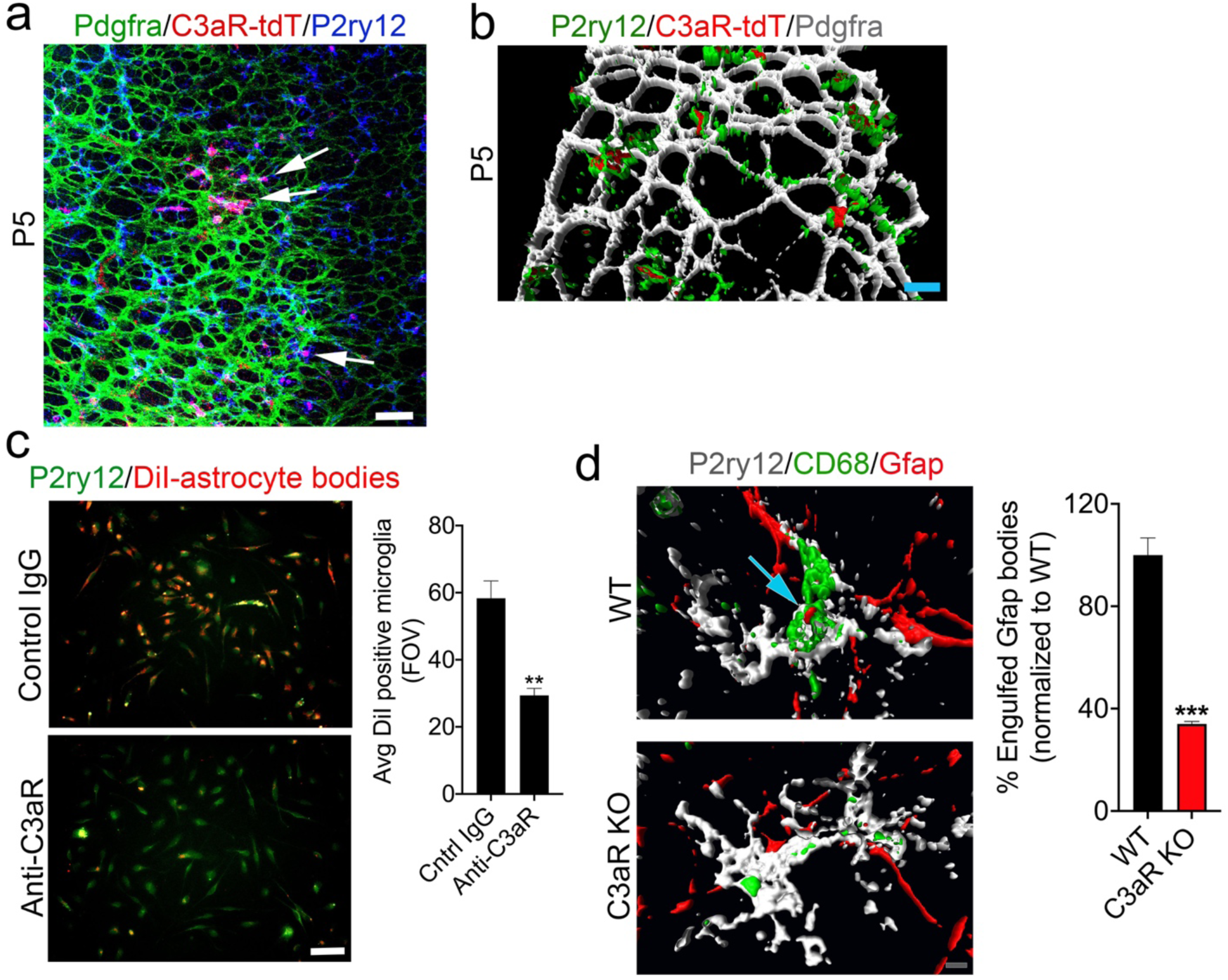
Microglia express C3aR and loss of C3aR reduces microglial pruning of astrocytes. **a.** Representative P5 C3aR-tdTomato (C3aR-tdT) expressing retinal flatmount showing Pdgfra and P2ry12 immunolocalization around the vascular growth front (n=3). Arrows indicate the localization of tdTomato in P2ry12-expressing microglia. **b.** Representative 3D-reconstructed image from C3aR-tdTomato expressing P5 retinal flatmount immunostained for P2ry12 and Pdgfra revealing C3aR reporter expression in microglia (P2ry12, green) that are closely interacting with astrocytes (Pdgfra, grey) (n=3). **c.** Primary microglial cultures isolated from P5 retinas were treated with DiI-labeled retinal astrocyte apoptotic bodies in the presence of control IgG or anti-C3aR IgG for 2 hours, then the number of DiI-astrocyte bodies engulfed by microglia in the two treatment conditions were quantified and shown in the bar graph (n=3). **d.** Z-stack images were acquired from P5 WT and C3aR KO retinal flatmounts immunostained for P2ry12, Gfap, and CD68 and 3D-reconstructed to visualize microglia (P2ry12) engulfed astrocyte (Gfap) bodies within microglial lysosomal/endosomal membrane protein CD68. Bar graph shows the percentage of Gfap debris localized within P2ry12, CD68 co-labelled microglia between WT and C3aR KO retinas (n=3). All error bars represent ± S.E.M. Statistical differences were calculated by unpaired *t*-test. ** P < 0.01, *** P < 0.001, and **** P < 0.001. Scale Bars: **a** is 75 µm, **b** and **d** are 10 µm, **c** is 40 µm.

**Figure 6.**
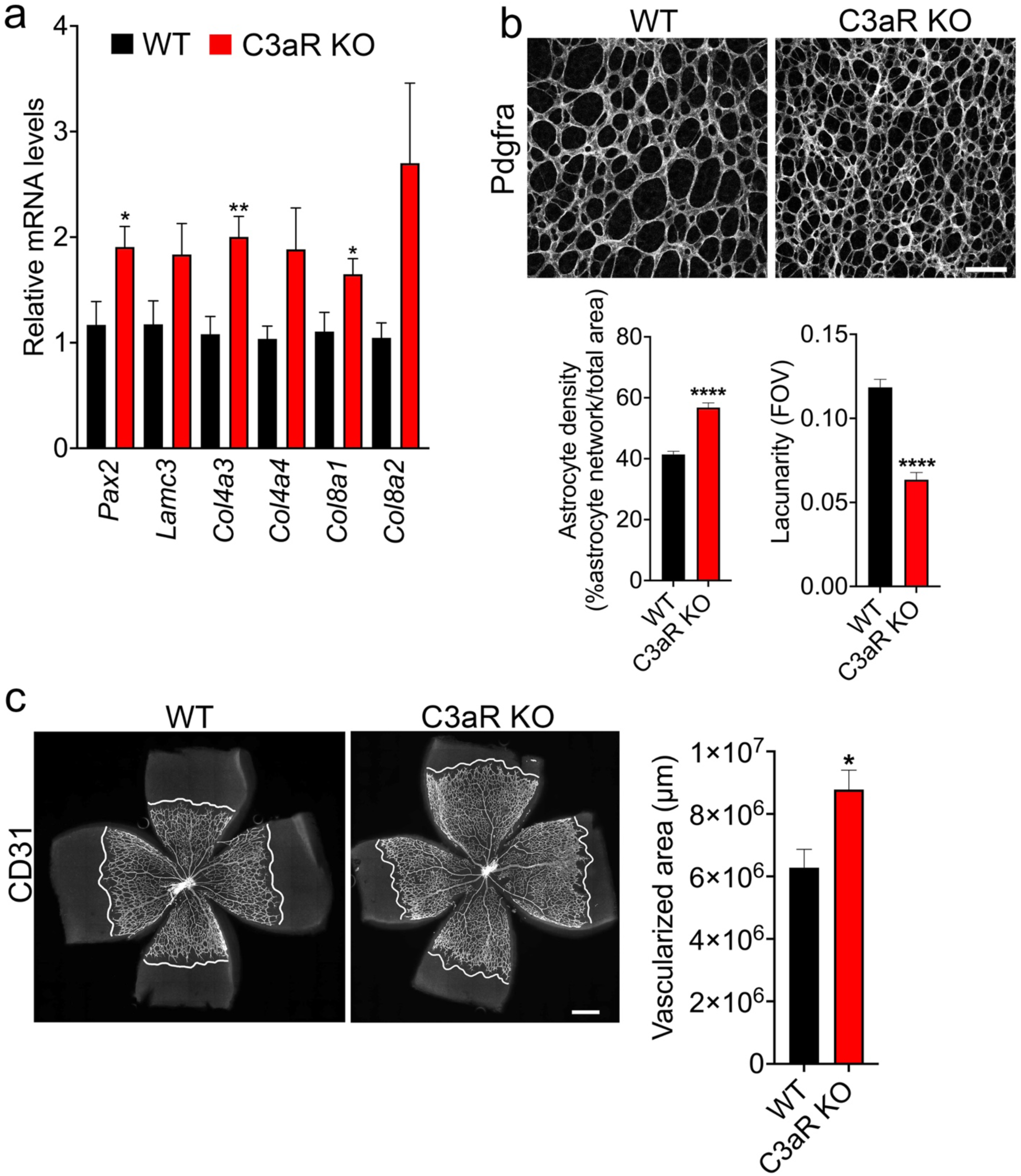
Loss of C3aR increases vascular growth rate and density at P5. **a.** Real-time PCR data showing gene expression levels of *Pax2*, *Lamc3*, *Col4a3*, *Col4a4*, *Col8a1*, and *Col8a2* in WT and C3aR KO P5 retinas (n=6). **b.** Representative P5 retinal flatmounts from WT and C3aR KO immunostained for Pdgfra showing astrocyte template patterning in the avascular region (n=5). Bar graphs represent the quantitative measurements of astrocyte density and lacunarity using ImageJ AngioTool software (n=5). **c.** Representative P5 Retinal flatmounts of WT and C3aR KO immunostained for CD31 showing vascular outgrowth and quantification of vascularized area in WT and C3aR KO (n=5). Scale bars: **b** is 50 µm and **c** is 500 µm. All error bars represent ± S.E.M. Statistical differences between WT and C3aR KO was calculated by unpaired *t*-test. * P < 0.05, ** P < 0.01, and **** P < 0.0001.

Moreover, subsequent superficial vascular growth and vascular density were increased with reduced spatial vascular patterning in P5 C3aR KO retinal flatmounts compared to WT littermate controls (**Fig. 6c and 7a)**. At P15 (retinal vascularization is near complete stage), C3 deficient retinas did not show significant defects in astrocyte or vascular patterning (data not shown). While, C3aR deficient retinas displayed areas of abnormal astrocyte morphology, the presence of non-vascular associated CD31^+^ endothelial cells, and areas of reduced microglial interaction with astrocytes and the superficial vasculature (**Fig. 7b**). These results illustrate that microglial C3aR is necessary for spatial patterning of astrocytes and organized succeeding superficial vascular network formation.

**Figure 7.**
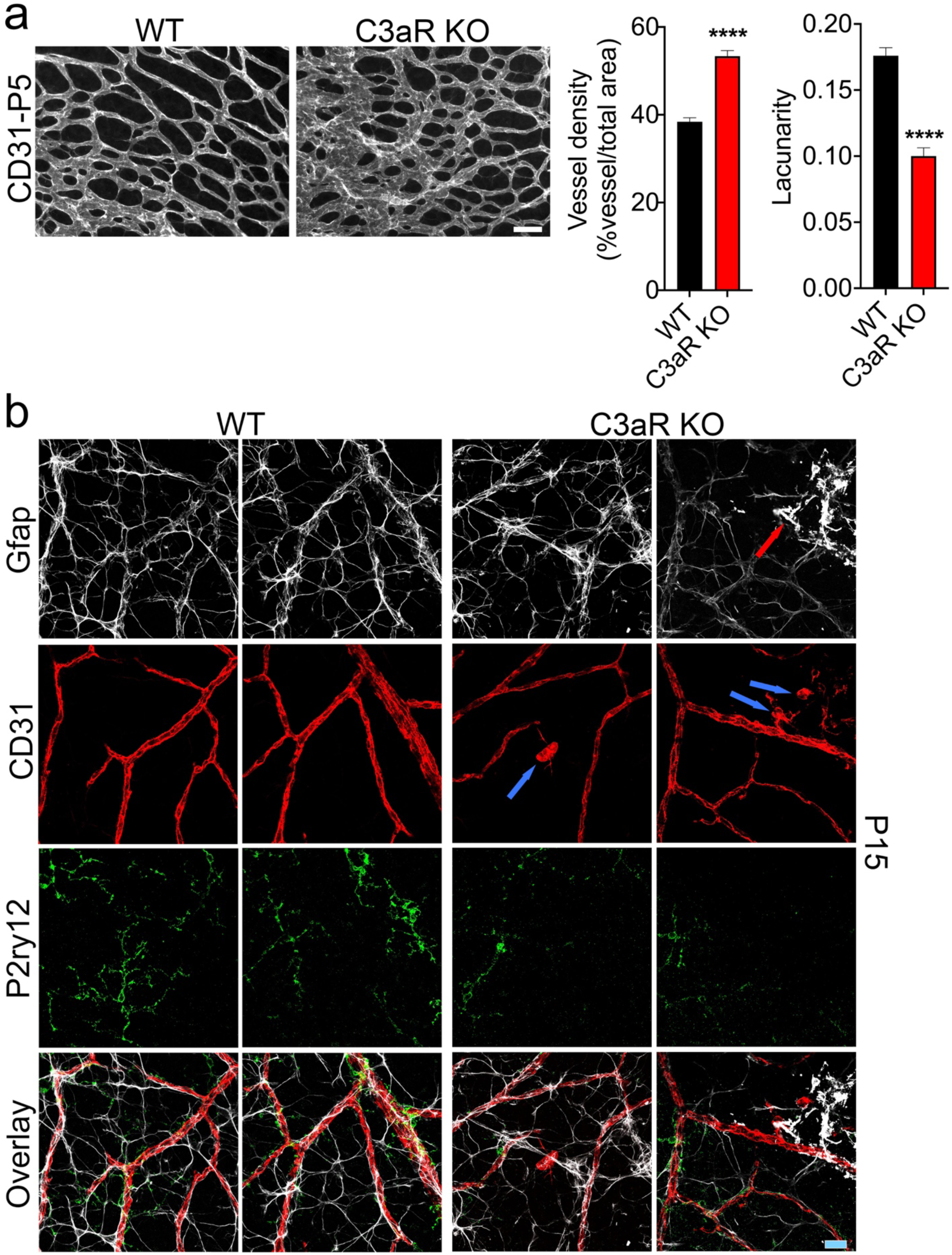
Loss of C3aR results in increased vascular density at P5 and defective astrocyte, microglial, and endothelial interactions at P15. **a.** P5 retinal flatmounts of WT and C3aR KO were immunostained for CD31 and images were taken in the vascularized central retina and the differences in vessel density and spatial branching (lacunarity) were measured and quantified using ‘ImageJ AngioTool’ software tool (n=5). **b.** Representative P15 retinal flatmounts of WT and C3aR KO immunostained for CD31, Gfap, and P2ry12 revealing endothelial, astrocytes, and microglial association (n=4). Red arrow points to dysmorphic Gfap immunostained astrocytes and blue arrows point to CD31 immunostained dysmorphic endothelial cells. Scale bar: 20 µm. All error bars represent ± S.E.M. Statistical differences between WT and C3aR KO was calculated by unpaired *t*-test. **** P < 0.0001.

In summary, these results identify microglia as a key regulator of astrocyte spatial patterning and subsequent vascular development through the C3a/C3aR derived signaling axis.

## Discussion

Microglia are essential for neural development [22,21,24], and in particular, they play an important role during neurovascular growth [4, 5]. In this study, we uncovered a mechanism through which microglia direct neurovascular development in the retina. Results from this study show that the depletion of microglia early during astrocyte patterning and subsequent primary superficial vascular growth phases increased astrocyte density and caused clustering of astrocyte template. Mechanistically, our study identified that C3/C3aR activation in microglia regulates spatial patterning of astrocyte and vascular networks. Specifically, deficiency of C3 resulted in a dramatic reduction of microglial phagocytosis of astrocyte debris leading to increased astrocyte density and reduced spatial gap. Furthermore, loss of C3 resulted in the upregulation of ECM genes transcript levels that are linked to vascular development [52,14,53]. In addition, loss of C3aR (a binding receptor for C3a) diminished microglial phagocytic removal of astrocytes, leading to an increase in astrocyte patterning and subsequent vascular growth.

Recent advancements in the discovery of microglial specific markers such as P2ry12, which are highly expressed during development [41], allowed us to further define the extensively ramified morphology of microglia along with their close interactions with astrocytes during development. The finding of the elaborate ramified morphology of microglia association with astrocytes and endothelial cells during development is particularly intriguing, as prior use of generic myeloid cell markers such as Iba1 did not reveal the complex morphology or the extent to which microglia associate with astrocytes during development [4,46,5]. The identification of close microglial interactions with astrocytes during migration and patterning suggest a role for microglia in assisting astrocyte template assembly. It is known that the basement membrane molecule laminin, present in the inner limiting membrane (ILM), provides a substrate for astrocyte migration and template formation in the retina [16,14,17]. Deletion of laminin isoforms or the laminin-binding proteoglycans severely disrupts ILM formation, astrocyte template assembly, and succeeding vascularization [16,14,17]. This study further discovered that microglial depletion causes abnormal clustering of astrocytes leading to altered spatial patterning of astrocyte matrices. Thus, these results strongly suggest that the spatially organized honeycomb shaped astrocyte patterning is regulated at multiple cellular and extracellular levels [6,14,17,15], allowing for the formation of the subsequent superficial vascular network in the retina during development.

Spatial restructuring during development is an important process that helps retinal maturation and function [22,39,56–58]. It is well documented that during the developmental restructuring process, large numbers of retinal ganglion cells are eliminated by microglia in part through the activation of the complement system [22, 58]. Particularly, C3 and C1q are reported to facilitate the developmental neural restructuring process [22,59,24]. Similar to ganglion cells [22, 58], large numbers of astrocytes are also removed by microglia peaking at P5 through unknown mechanisms [39, 40]. Our study shows that C3 deficiency (but not C1q loss) resulted in increased astrocyte density, reduced astrocytes spatial gap, and increased vascular density at P5. Although we cannot rule out the involvement of other complement components, our data strongly suggest that C3 plays a substantial role in facilitating the spatial establishment of astrocyte matrices for subsequent superficial vascular development.

Interestingly, loss of CR3, a well-known receptor involved in phagocytic elimination of retinal ganglion cells [22], did not significantly alter astrocyte patterning or subsequent vascular growth. On the other hand, loss of C3aR (receptor for cleaved small C3a fragment) resulted in increased astrocyte and vascular density. C3a anaphylatoxin activation of C3aR is widely regarded as a function involved in an inflammatory response [29, 60]. Our data show that local microglial expression of C3 and its receptor C3aR play a crucial role during retinal vascular development. It is likely that microglial local deposition of C3 over the astrocytic template activates C3aR to prune and facilitate superficial vascular patterning. Supporting our notion, loss of C3 or C3aR dramatically reduced microglia’s phagocytic ability to eliminate astrocytes during retinal development, leading to increased astrocyte and vascular density.

Deposition and remodeling of the ECM is important for promoting vascular growth and stability [61, 62]. Astrocytes are known to secrete ECM proteins that provide a substrate for endothelial tip cell migration and the formation of superficial retinal vasculature [12,14,63,15]. Defect or changes in ECM molecular composition could alter vascular growth and vascular matrix integrity [64,14,65]. As prior report indicate a role for C3a-C3aR axis in regulating proteases mediated ECM remodeling [66], enrichment of ECM related genes in C3 and C3aR deficient retinas suggest a possible defect in remodeling of ECM components involved in angiogenesis [64,14,65]. Although further studies are required to determine C3/C3aR involvement in remodeling of ECM during retinal angiogenesis, our thus far findings provide a compelling role for complement in creating spatially organized astrocyte matrices to fine tune superficial retinal vascular morphogenesis.

Taken together, our study identified a novel functional role for the microglia-derived C3/C3aR axis in defining astrocyte spatial pattering for subsequent primary superficial retinal vascular growth during retinal development.

## Materials and Methods

### Mice

All animal procedures were performed in accordance with the Massachusetts Eye and Ear Animal Care Committee. Mice were maintained in a room with a 12 hours light/12 hours dark cycle. C57BL6/J (stock # 00664), C3 mutant (referred as KO) (stock # 029661), and C3aR mutant (referred as KO) (stock # 033904) breeding pairs were purchased from Jackson Laboratories. C3 KO and C3aR KO mice were crossed to C57BL6/J mice to create a heterozygous line. Pups generated from the heterozygous lines carrying C3+/+ (wild type (WT)), C3-/- (C3 KO), C3aR+/+, and C3aR-/- genotypes were used for the studies. Of note, it is reported that mutant C3 mRNA was detected in extrahepatic tissues of deficient mice with no detectable level of active cleaved C3 proteins [67, 68], our RNA-seq analysis also showed C3 mRNA expression in C3 mutant mice. Floxed C3 IRES-tdTomato reporter mice [69] obtained from Dr. Kemper were bred to C57BL6/J mice and maintained on a C57BL6/J background. Floxed C3aR-tdTomato reporter knock-in mice [55] obtained from Dr. Köhl were maintained on a C57BL6/J background.

### Microglial depletion during development

Timed-pregnant mice were maintained on a control chow diet or a chow diet containing Csfr1 inhibitor (PLX5622, Plexxikon Inc. Berkeley, CA) from gestational day 13.5/14.5. Retinas were collected from pups at postnatal days (P) 1, P5, and P15 for downstream analysis.

### Isolation and culture of retinal microglia and astrocytes

Astrocytes and microglia were isolated by immunopanning methodology [70, 71]. In brief, six-well culture plates were coated with their respective secondary antibodies (10µg/ml diluted in sterile 50mM Tris-HCl, pH 9.5) by incubating for 1 hour at 37^•^C. After washes, culture plates were coated with 2.5 µg/ml of anti-CD11b or anti-integrin beta5 overnight in the cell culture hood and washed in 1X PBS. P5 retinas (n=8) were isolated in ice-cold sterile 1X-DPBS (Thermo Fischer Scientific, Foster City, CA) and incubated in DMEM/F12 containing 5% fetal bovine serum (FBS), collagenase D (1mg/ml,) and DNase (0.1mg/ml) for 30 min at 37^•^C. The retinas were then gently triturated in DMEM/F12 supplemented with 10% FBS and filtered through a 40µm cell strainer to remove cell clumps and debris. The resultant single cell suspension was incubated in asecondary antibody coated well for 10 min at 37^•^C to deplete non-specific cell binding. The unbound cell suspension was then transferred to anti-CD11b coated wells and incubated for 20 min at 37^•^C to capture microglia (culture plate was shaken every 5 min to dislodge non-specific cell binding). After depleting microglia, the cell suspension was transferred to integrin beta 5 coated wells to capture astrocytes and incubated for 45 min at 37^•^C (culture plates were shaken every 10 min to dislodge non-specific cell binding). After removing the unbound cells, microglial and astrocyte cell culture plates were thoroughly washed in serum free DMEM/F12. Microglial cells were grown and maintained in DMEM/F12 supplemented with 5% FBS and 10ng/ml of recombinant mouse csf1, and astrocytes were grown and maintained in DMEM/F12 supplemented with 10% FBS and N-2.

### In vitro phagocytosis assay

To examine microglial phagocytosis of dead astrocyte bodies, ∼2,500 microglial cells were plated on a 12mm glass coverslip and maintained for 48 hours. Astrocyte cell suspensions were labeled with Vybrant™ Dil cell labeling solution as suggested by the manufacturer (Thermo Fischer Scientific, Foster City, CA) and cell death was induced by incubating the labeled astrocyte cell suspension with 1µM staurosporine (Cayman Chemical Company, Ann Arbor, MI). Microglial cells were then treated with DiI-labeled dead astrocyte bodies in 1:1 ratio and incubated for 2 hours. For receptor neutralization assays, microglial cells were preincubated with 20µg/ml of anti-C3aR antibody (Catalog # MAB10417) or control IgG (R&D Systems, Inc. Minneapolis, MN) suspended in DMEM/F12 supplemented with 5% heat-inactivated FBS, followed by treatment with DiI-labeled astrocyte dead bodies in the presence of antibodies for 2 hr at 37^•^C. After thorough washes, cells were fixed in 4% PFA, blocked for 1hr at room temperature (in 10% fetal bovine serum, 0.05%triton-X100, and 0.01% sodium azide in 1X PBS), followed by incubation with P2ry12 antibody (Catalog# AS-55043A, AnaSpec, Fremont, CA) overnight at 4^•^C and then incubated with goat anti-rabbit Alexa Fluor 488 or 647 (Thermo Fischer Scientific, Foster City, CA), images were acquired using epifluroscent microscope (Axio Observer Zeiss).

### Flatmount preparation and immunostaining

Retinal flatmounts were prepared as described previously [14]. In brief, eyes were fixed in 4% paraformaldehyde (PFA) for 10 min and then dissected in 1X PBS. After discarding the anterior chamber and lens, the retina was separated from the sclera. Retinal flatmounts were prepared with four radial incisions and the flatmounts were then stored in methanol at -20^•^C until immunostaining was performed. For immunohistochemistry, retinas were rehydrated in 1X PBS and washed at 3X and blocked in blocking buffer (10% fetal bovine serum, 0.05%triton-X100, and 0.01% sodium azide in 1X PBS) for 2hr at room temperature. Following blocking, retinas were then incubated with primary antibodies for 24 hr at 4^•^C. After washes in 1X PBS, retinal flatmounts were incubated with respective secondary antibodies for 4hr at room temperature or overnight at 4^•^C. The flatmounts were then washed in 1X PBS and mounted onto a slide using anti-fade medium (Permaflour, Thermo Fisher Scientific, Waltham, MA).

Primary antibodies used: Rat anti-Pdgfra (CD140A, clone: APA5, BD Biosciences, San Jose, CA), anti-chicken GFAP (Aves Labs, Inc, Davis, CA), rabbit anti-P2ry12 (a gift from Dr.Butovsky, Brigham and Women’s Hospital), rat anti-mouse CD68 (Clone: FA-11, Biolegend, Dedham, MA), anti-rabbit Iba1(Catalog# 019-19741, FUJIFILM Wako Chemicals, Richmond, VA), and goat anti-CD31 (Catalog # AF3628, R&D Systems, Inc. Minneapolis, MN).

Secondary antibodies used: Donkey anti-chicken-594 (Biotium, Inc. Fremont, CA), donkey anti-rat-594, donkey anti-rabbit-488, and donkey anti-goat-647 (Thermo Fisher Scientific, Waltham, MA)

### Cryosectioning

The enucleated eyes were fixed in 4% PFA for 2hr at room temperature and then the dissected posterior eyecups were cryopreserved in 10%, 20%, and 30% sucrose and then frozen in Tissue-Tek® O.C.T compound (Ted Pella, Inc. Redding, CA). The eyecups were then cut at 12µm thickness and used for immunohistochemistry.

### Flow cytometry

Singe cell suspensions were prepared from retinas that were dissected in ice-cold HBSS and then incubated with collagenase D (1mg/ml) and DNAse (0.1mg/ml) (Millipore Sigma) in DMEM/F12 supplemented with 5% FBS for 30 min at 37^•^C. After filtering the cells through a 40µm cell strainer, single cell suspensions were blocked with anti-mouse CD16/32 monoclonal antibody and then stained with APC anti-Pdgfra (CD140a, clone: APA5), and PE anti-p2ry12 (clone: S16007D, Biolegend, Dedham, MA). Dead cells were distinguished with DAPI stain. The data for flow cytometry was acquired by Cytoflex S (Beckman Coulter, Indianapolis, Indiana) and analyzed using FlowJo version 10.1.

### Real-time PCR

Retinas were isolated in ice-cold PBS and RNA was extracted using RNA STAT-60™ as recommended by the manufacturer. After determining the RNA concentration using NanoDrop™, 500ng of total RNA was used for cDNA synthesis using SuperScript™ IV VILO™ Master Mix. Real-time PCR reactions were performed in the CFX384™ Real-time PCR platform (Bio-Rad, Hercules, CA) using SYBR Green master mix (Applied biosystems, Thermo Fischer Scientific, Foster City, CA). to determine the relative expression level of Pdgfa, P2ry12, Tmem119, Csf1r, Vegfc, Vegf120, Vegf164, and Vegf188 [72]. Sequences of primers used: Ppia (TTC ACC TTC CCA AAG ACC AC and CAA ACA CAA ACG GTT CCC AG), Pdgfa (ATTAACCATGTGCCCGAGAA and GTATCTCGTAAATGACCGTCCTG), P2ry12 (CCAGTCTGCAAGTTCCACTAAC and GAGAAGGTGGTATTGGCTGAG, Tmem119 (GGTCCTTCACCCAGAGC and GGAGTGACACAGAGTAGGC), Csf1r (TGT ATGTCTGTCATGTCTCTGC and AGGTGTAGCTATTGCCTTCG), Vegfc (GAAGTTCCACCATCAAACATGC and CAGCGGCAT ACTTCTTCACTA), Vegf120 (GCCAGCACATAGGAGAGATGAGC and CGGCTTGTCACATTTTTCTGG), Vegf164 (GCCAGCACATAGGAGAGATGAGC and CAAGGCTCACAGTGATTTTCTGG), and Vegf188 (GCCAGCACATAGGAGAGATGAGC and AACAAGGCTCACAGTGAACGCT), B2m (TGG TCT TTC TGG TGC TTG TC and GGG TGG AAC TGT GTT ACG TAG), Col8a1(TTGCCTAGCAACTGGGTGAG and TATGAGCTCCTGTGGGCTCT), Col8a2 (CCCCGGTAAAGTATGTGCAG and GGCTCTCCTTTCAAGTCCATT), Col4a4 (GTTGTACATGGAAGGACAGGAG and ACTTGGTGGATGTTGCAGTAG), Col4a3 (GAACTGTAGGGGACATGGGC and CTGGATCACCTCTGACACCG). Gfap (AACCGCATCACCATTCCTG and GCATCTCCACAGTCTTTACCA), Pax2 (TTCCTATCCCGATGTCTTCCA and GATGCAGATAGACTGGACTTGAC), Lamc3 (TGCTAGCAGCAGCTGAACAT and TCAGTGGGATAGGACAGGGG).

### mRNA sequencing

RNA was extracted as described above from P5 WT and C3 deficient retinas and the integrity of RNA was assessed by Bioanalyzer (Agilent 2100). RIN values for all the samples used for downstream cDNA library construction were >9.0. To determine global gene expressional changes, mRNA was purified with oligo d-T attached magnetic beads (NEBNext®) and 500 ng of RNA was used for cDNA library construction with NEBNext ultra II directional RNA library prep kit for Illumina as recommended by the manufacturer (New England Biolabs, Inc, Ipswich, MA). cDNA libraries were validated on the TapeStation (Agilent) and the sequencing was performed with Illumina Nextseq 2000 platform with a target of 25million reads per sample.

### RNA-seq analysis of differential gene expression pattern

Transcriptome mapping was performed using the STAR aligner [73] and the mm9 assembly of the mouse reference genome. Read counts for individual transcripts were obtained using HTSeq [74] and the GENCODE M1 (NCBIM37) gene annotation. Differential expression analysis was performed using the EdgeR package [75] after normalizing read counts and including only genes with CPM > 1 for at least one sample. Differentially expressed genes were defined based on the criteria of >1.5-fold change in normalized expression value and false discovery rate (FDR) <0.05. Heatmaps and PCA plots were generated using normalized gene expression values (log2 FPKM). Heatmaps were generated using the R package pheatmap [76]. Analysis of enriched functional categories among detected genes was performed using EnrichR [51].

### Image analysis

Samples were imaged using an epifluroscent microscope (Axio Observer Zeiss) or confocal microscopy (SP8, Leica). For 3D image reconstruction of Z-stack images, Amira® 2019.4 software tool was used.

### Analysis of astrocyte and vascular coverage

Tiled images of the entire retinal surface immunostained for Pdgfra or CD31 were acquired. Using NIH-ImageJ (version 2.1.0/1.55c)) software, scale measurements were set and an outline of the astrocyte or vascular growth front was created using the manual freehand selection tool to measure the total area immunostained by Pdgfra or CD31. Tip cells were manually quantified in all four quadrants using ImageJ multi-point tool.

### Microglia immunostained cellular area measurement

Images of retinal flatmounts co-labeled with CD31 and P2ry12 or CD31 and Iba1 were acquired in all the retinal quadrants near the tip cell and avascular peripheral regions by confocal microscopy. The outline of the P2ry12 or Iba1 stained cell area was manually drawn using Image J (version 2.1.0/1.53c) freehand selection tool and each immunostained cell area was measured and quantified.

### Analysis of astrocyte and vessel density, branching, and lacunarity

The astrocyte and blood vessel covered areas were quantified using ImageJ (version 2.1.0/1.53c) Angiogenesis Analyzer plugin tool. In brief, images stained for Pdgfra or CD31 were opened via AngioTool, scale measurement was entered, and data measurements containing vascular density, branching index, and lacunacity that were automatically generated.

### Statistical analysis

All the other graphs and statistical analysis were performed using Prism version 9 software. All data were presented as the mean +/-SEM. Statistical differences between the two groups were determined by an unpaired t-test. The ‘n’ and the level of statistical significance were noted in the legend of each figure.

### Data availability

RNA-seq data was upload to the GEO database repository. Images and raw flow cytometry data used for the analysis are uploaded to Harvard institutional repository. (https://dataverse.harvard.edu/dataset.xhtml?persistentId=doi:10.7910/DVN/CDJYDT). Upon acceptance these data will become publicly available. All raw data will also be available by reaching out to the Lead Contact, Gopalan Gnanaguru.

### Author contributions

GG and KMC designed research and wrote the manuscript; GG performed research and analyzed data; SJT and KY analyzed data, RS and GMB assisted with RNA-seq studies and data analysis, JK generated and provided reporter mice.

## Acknowledgments

This work was supported by NIH/National Eye Institute Grant R01EY032502 (to G.G) and NIH/NIDDK P30 DK040561 (to R.I.S). We thank Plexxikon, inc, for providing PLX5622 chow diet. We sincerely thank Dr.Butovsky (Department of Neurology, Brigham and Women’s Hospital, Harvard Medical School) for providing the P2ry12 antibody and Dr. Claudia Kemper (Immunology Center, National Heart, Lung, and Blood Institute, NIH, Bethesda, MD, USA 20892) for providing the Floxed C3 IRES-tdTomato reporter mice. We thank MGH Nextgen sequencing core staff Danning Zhou and Ulandt Kim for the help with cDNA library construction and sequencing.

## Conflict of interest

The authors declare no competing interests.

## Notes

### Competing Interest Statement

The authors have declared no competing interest.

### Summary of Updates

This manuscript has been revised to update Figure1, Figure 2, figure 3, Figure 4, Figure 5, Figure 6, and Figure 7

