## Supplementary data for "Complement facilitates developmental microglial pruning of astrocyte and vascular networks"

**Supplementary figures**

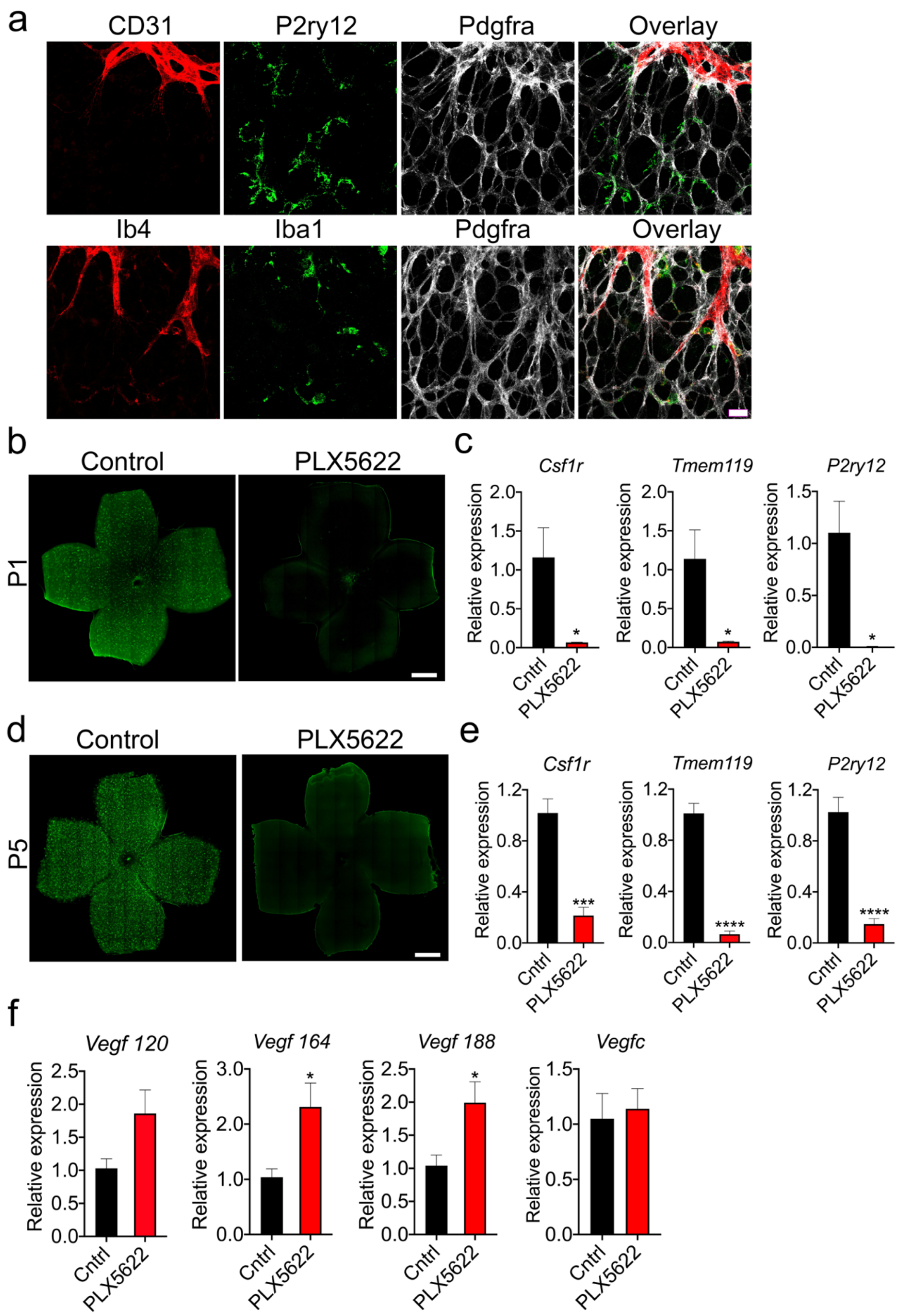

**Supplementary fig. 1**. Csf1r specific antagonist effectively depletes retinal microglia early in the vascular developmental stage. **a**) Top panel: Representative P5 retinal flatmount showing immunostaining of CD31 (endothelial cell marker), P2ry12 (microglial marker), Pdgfra (astrocyte marker), and overlaid image around vascular growth front (n=5). Bottom panel: Representative P5 retinal flatmount showing immunostaining of isolectin B4 (endothelial and immune cell marker), Iba1 (microglial/macrophage marker), Pdgfra (astrocyte marker), and overlaid image around vascular growth front (n=5). (**b-e**) Time-pregnant C57BL/6J mice from gestational day E13.5/E14.5 were fed a control or Csf1r specific antagonist (PLX5622) diet and the litters were analyzed at P1 and P5 developmental time points. (A-F). **b)** Representative P1 retinal flatmounts from control or PLX5622 treated groups showing staining of P2ry12 (Green). **c)** Real-time PCR analysis of relative gene expression levels of microglia markers, *Csf1r*, *P2ry12*, and *Tmem119* at P1 (n=3) in the retinas of control or PLX5622 groups. **d)** Representative anti-P2ry12 immunostaining of retinal flatmounts showing microglial distribution at P5. **e)** Real-time PCR analysis showing relative gene expression levels of microglial markers, *Csf1r*, *P2ry12*, and *Tmem119* at P5 (n=5) in the retinas of control or PLX5622 groups. **f)** Real-time PCR analysis showing relative gene expression levels of *Vegfa* isoforms (120, 164, and 188) and *Vegfc* at P5 (n=5) in the retinas of control or PLX5622 groups. Scale bar- **a** is 15μm, **b** and **d** are 500μm. All error bars represent ± S.E.M. Statistical differences between control and PLX-5622 group was calculated by unpaired *t*-test. * $P < 0.05$, *** $P < 0.001$, **** $P < 0.0001$

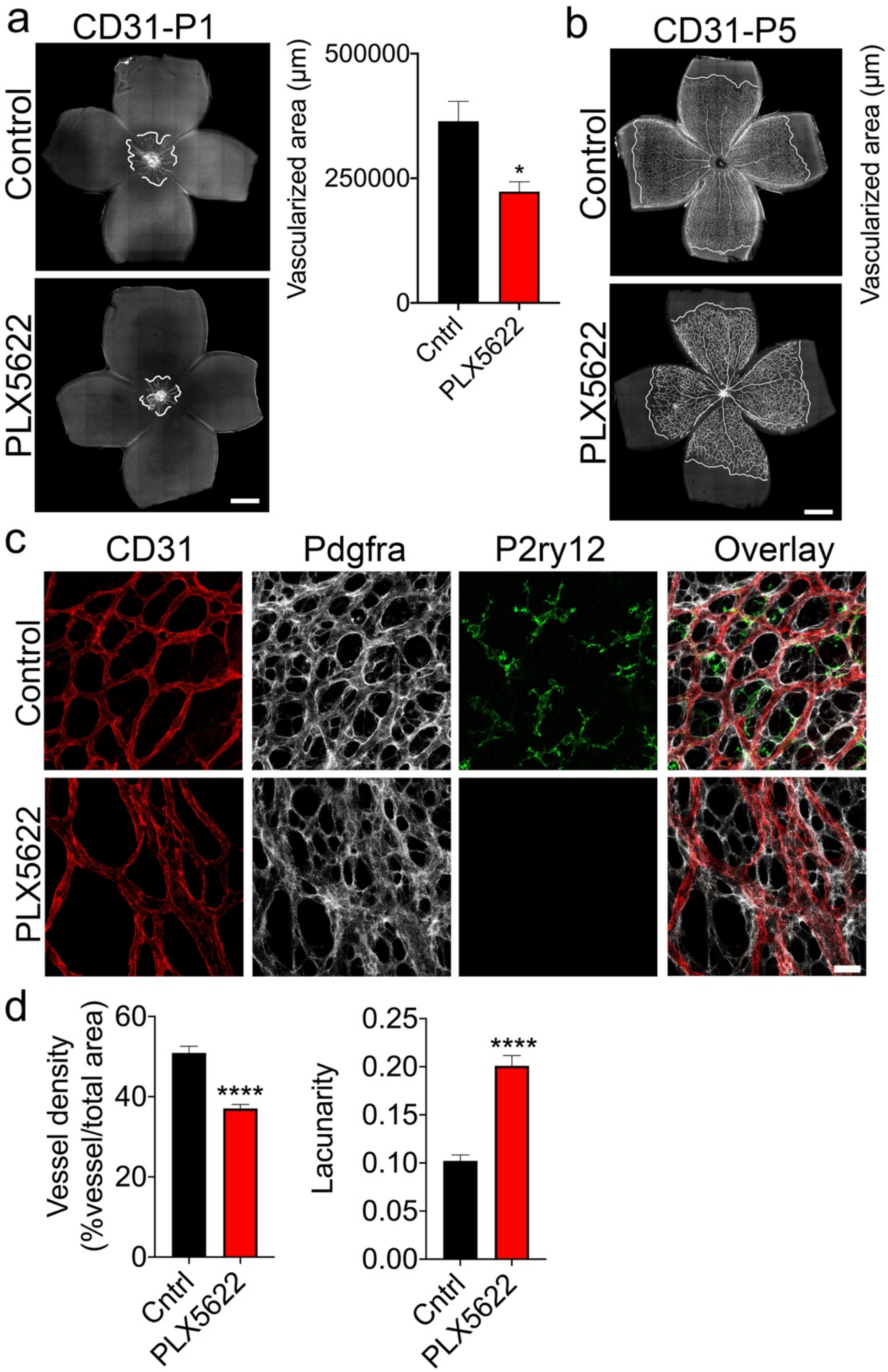


**Supplementary fig. 2**. Microglial depletion reduces vascular growth and density. Time-pregnant C57BL/6J mice from gestational day E13.5/E14.5 were fed a control diet or diet containing Csfr1 specific antagonist (PLX5622) and the littermates were analyzed at respective ages (a-d). **a and b)** P1 and P5 retinal flatmounts from control and PLX5622 group immunostained for CD31 (endothelial cell marker). The vascularized areas (indicated by dotted lines in the images) were quantified and shown in the bar graphs (n=4). **c)** Representative images of P5 retinal flatmounts from control and PLX5622 group immunostained for CD31, Pdgfra, and P2ry12 showing microglial interaction with endothelial cells and astrocytes in the vascularized area (n=5). **d)** Images of control and PLX5622 P5 retinal flatmounts immunostained for CD31 in the vascularized central region of all four quadrants were acquired. Using 'NIH-ImageJ AngioTool' software, vessel density and lacunarity were quantified and shown (n=5). Scale bars: **a** and **b** are 500 µm, **c** is 50µm. All error bars represent ± S.E.M. Statistical differences between control and PLX5622 group were calculated by unpaired *t*-test. * $P < 0.05$, ** $P < 0.01$, *** $P < 0.001$, **** $P < 0.0001$

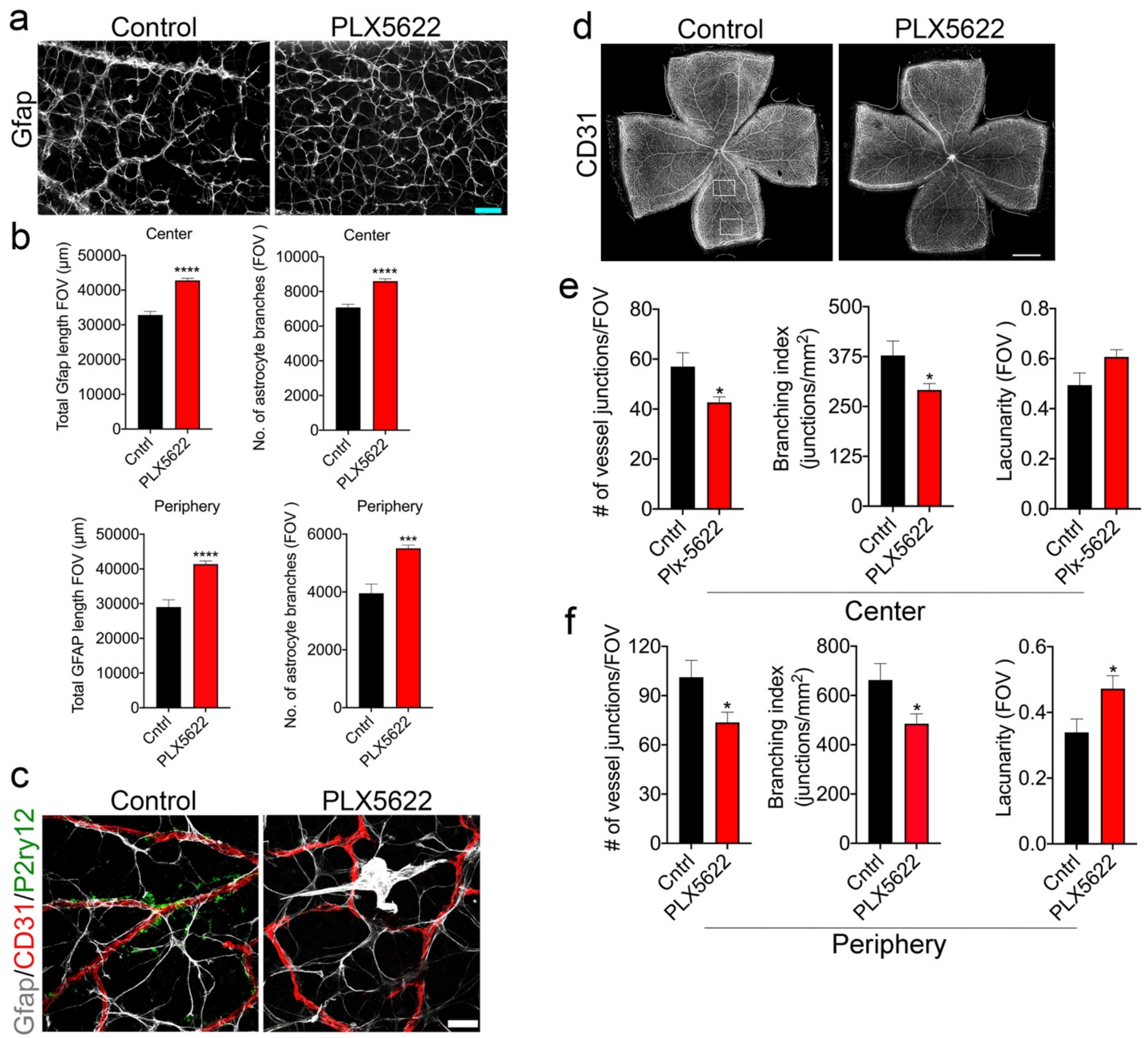


**Supplementary Fig. 3.** Increased astrocyte patterning and reduced vascular density persists in microglia-depleted retinas at P15. **a)** Representative P15 retinal flatmounts from control and PLX5622 group immunostained for Gfap showing astrocyte patterning (n=3). **b)** Gfap immunostained images of the central and peripheral retina (representative image in a) were processed using ImageJ software to calculate the total Gfap immunostained length and branches (as data shown in c). **c)** Representative confocal image of P15 retinal flatmounts immunostained for Gfap, CD31, and P2ry12 showing astrocyte (Gfap-grey), microglia (P2r1y2-green), and blood vessel (CD31-red) morphology and distribution. **d)** Representative P15 retinal flatmount of control and PLX5622 group showing blood vessel patterning. Images were taken in the central and peripheral areas of all four quadrants (example of images acquired in the central and peripheral retina are shown as boxes) and further assessed using NIH-Angiogenesis Analyzer software to quantify vessel density and distribution. **e)** Quantification of CD31 immunostained images indicating number of vessel junctions, branching index, and lacunarity in the central region of P15 retina of control and PLX5622 groups. **f)** Quantification of CD31 immunostained images showing number of vessel junctions, branching index, and lacunarity in the peripheral region of P15 retina of control and PLX5622 groups. Scale bar: **a** is 50 µm, **c** is 20 µm, and **d** is 500 µm. All error bars represent ± S.E.M. Statistical differences between control and PLX-5622 group was calculated by unpaired *t*-test. * $P < 0.05$, *** $P < 0.001$, **** $P < 0.0001$

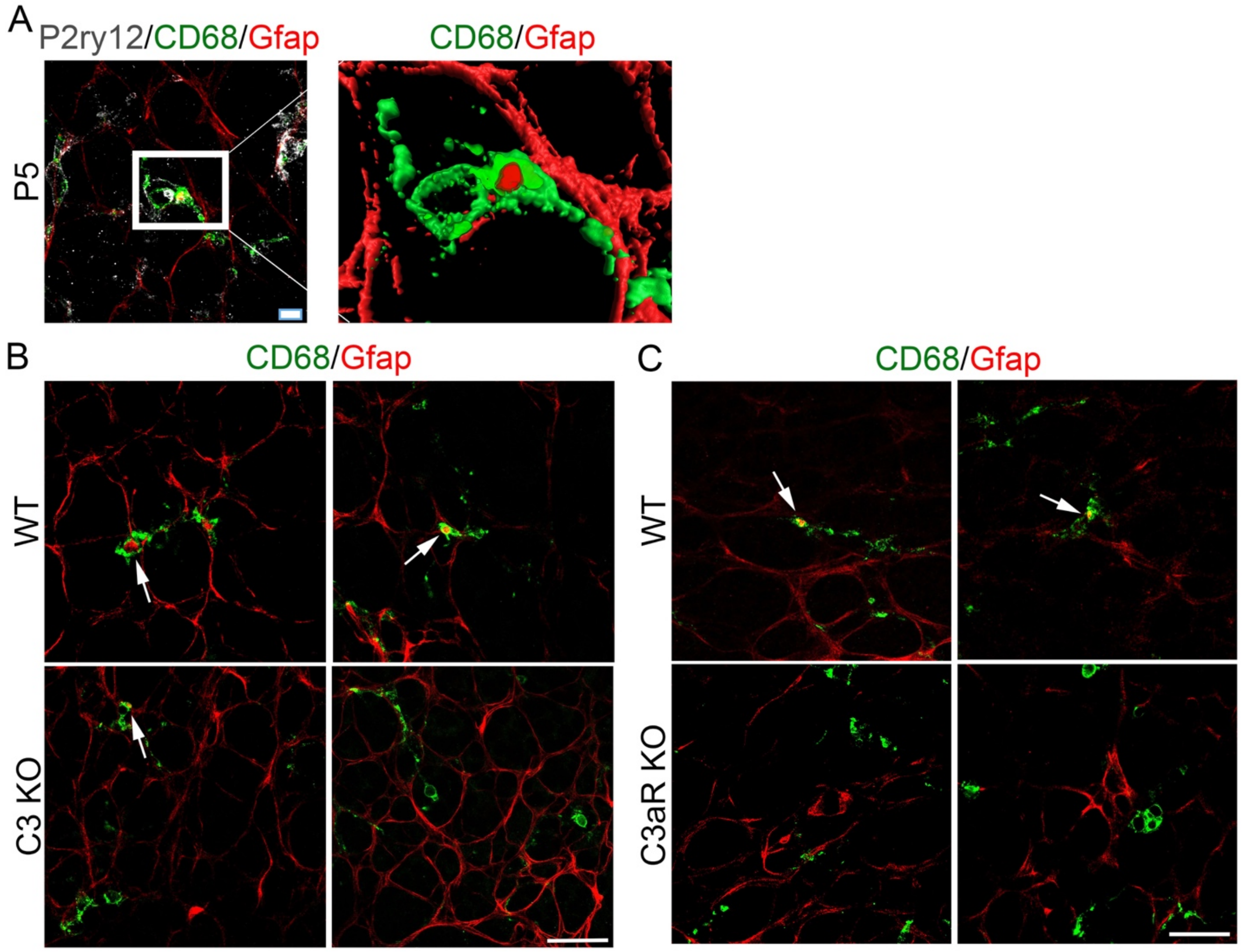


**Supplementary Fig. 4.** Loss of C3 and C3aR reduces microglial phagocytosis of astrocytes during template rearrangement. **a)** Left panel: Representative z-stack images of P5 retinal flatmount showing GFAP, P2ry12, and CD68 immunostaining (n=5). Right panel: Enlarged 3D-reconstructed boxed area in **(a)** that excludes microglial P2ry12 to better visualize internalized GFAP astrocyte debris within CD68+ lysosomes (green). **b)** P5 retinal flatmounts of WT and C3 KO were immunostained for CD68 and GFAP (n=3). High-magnified Z-stack images were acquired using confocal microscopy and Z-stack images were scanned through to reveal Gfap localization within CD68 immunostained microglia. **c)** P5 retinal flatmounts of WT and C3aR KO were immunostained for CD68 and GFAP (n=3). High-magnified Z-stack images were acquired using confocal microscopy and Z-stack images were scanned through to reveal Gfap localization within CD68 immunostained microglia. Scale bar: **a** is 10 μm, **b** and **c** are 15 μm.

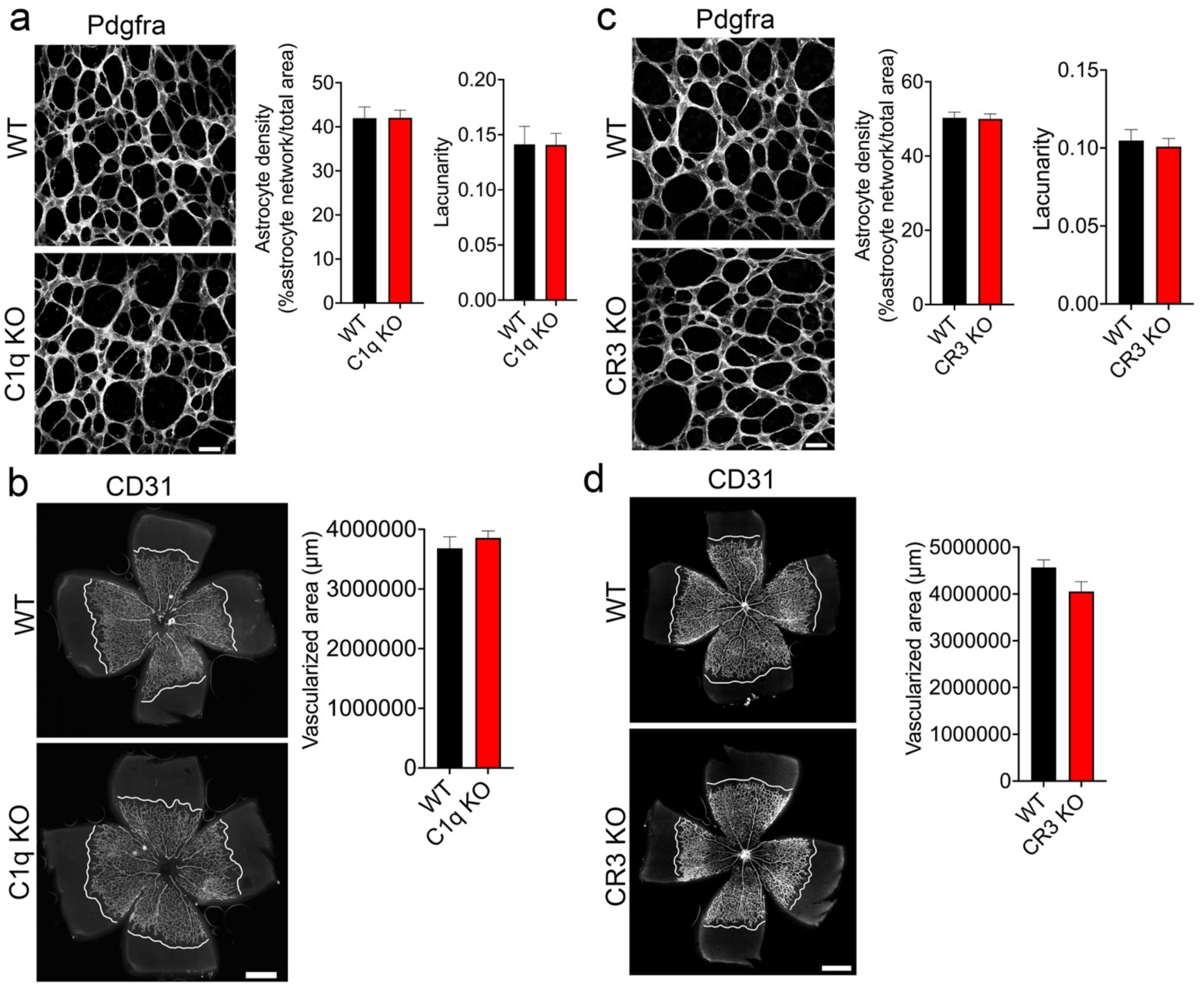


**Supplementary Fig. 5.** Loss of C1q or CR3 did not alter astrocyte patterning or vascular growth at P5. **a)** Representative P5 retinal flatmounts from WT and C1q KO immunostained for Pdgfra showing astrocyte template patterning in the avascular region (n=3). Bar graphs represent the quantitative measurements of astrocyte density, branching index, and lacunarity using ImageJ AngioTool software (n=3). **b)** Representative P5 retinal flatmounts of WT and C1q KO immunostained for CD31 displaying vascular outgrowth (indicated by dotted lines in the images). Differences in vessel density and spatial branching (lacunarity) were quantified in WT and C1q KO (n=3) using ‘ImageJ Angiogenesis Analyzer’ software tool. **c)** Representative P5 retinal flatmounts from WT and CR3 KO immunostained for Pdgfra showing astrocyte template patterning in the avascular region (n=4). Bar graphs represent the quantitative measurements of astrocyte density, branching index, and lacunarity using ImageJ AngioTool software (n=4). **d)** Representative P5 retinal flatmounts of WT and CR3 KO immunostained for CD31 displaying vascular outgrowth (indicated by dotted lines in the images). Differences in vessel density and spatial branching (lacunarity) were quantified in WT and CR3 KO (n=5) using ‘ImageJ Angiogenesis Analyzer’ software tool. Scale bar: **a** and **c** are 50 μm, **b** and **d** are 500 μm.
